# Sequence-Derived Representations versus Pfam-Domain Content for Biosynthetic Gene Cluster Retrieval

**DOI:** 10.64898/2026.08.21.746127

**Authors:** Rustambek Urokov, Asifullah Khan, Farhod Eshboyev, Dovud Asadov, Sejuti Rahman, Dilafruz Kushokova

**Author notes:** {, }.

## Abstract

Retrieving BGCs related to those of a known producer can be regarded as a representation-learning objective. We hypothesize that ESM-2 sequence-derived representations of BGCs can improve retrieval beyond the Pfam-domain content metric. Our toolkit is the following: group-disjoint train, validation, and test assignments, validation-frozen model selection, five seeds, and family-level paired inference. Of 6,953 atlas BGCs from 182 deduplicated *Streptomyces griseus* genome accessions, 5,325 silver-labeled BGCs are split into 98 training, 21 validation, and 21 test reference groups. Of the test reference groups, 16 are eligible for retrieval diagnostics. Pfam Jaccard scored Recall@50 of 0.8788, while Pfam-augmented BGC-SetNet scored 0.8472. The combination of ESM and Pfam-augmented BGC-SetNet scored 0.8769. A weighted Pfam Jaccard obtained a slightly higher score of 0.8789, which has a negligible difference compared to unweighted Pfam Jaccard. Our results do not support the claim that sequence-derived representations can recover alternative biosynthetic pathways on this benchmark. Instead, explicit Pfam remains the major signal for this objective. Our results define the curation and pathway-level validation processes that are necessary for a more robust biological test.

## 1 Introduction

Biosynthetic gene clusters (BGCs) consist of co-localized genes encoding enzymes which can be involved in biosynthesis of specialized metabolites. Given a known producer, retrieval systems should prioritize candidate BGCs that share relevant biosynthetic information, including divergent gene content. While these candidates are suitable for further investigation, computational similarity alone does not necessarily imply that they produce the same product through a different pathway.

Current domain-content methods include strong BGC-comparison baselines. While BiG-SCAPE combines domain-content similarity, domain duplication, and adjacency information [2], antiSMASH Known-ClusterBlast uses domain information [1]. We use maximum Pfam Jaccard and treat it as a simple Pfam-content baseline. Protein language models (PLMs) provide sequence-derived representations. ESM-2 representations are generated from protein sequences [5], but should not be treated as structural measurements.

We evaluate Pfam-augmented BGC-SetNet with a fixed group-disjoint split and then test two strategies to improve the representation: feeding the model with BGC-level Pfam inventory used by the baseline and learning weights for Pfam domains.

Our contributions are:

1. a reproducible group-disjoint framework for evaluating BGC retrieval that specifies unknown candidates, reference construction, and family-level inference;
2. a comparison of frozen ESM-2 representations, attention-based aggregation, and explicit Pfam-domain content;
3. a differentiable weighted-Pfam Jaccard model trained only on labels from the training families; and
4. an empirical analysis of the benchmark, illustrating that explicit Pfam-domain content performs better than the sequence-derived representations. The ESM plus Pfam-augmented BGC-SetNet ensemble shows small, statistically unsupported gains in several secondary metrics.

## 2 Related Work

### BGC comparison

For BGC comparison, antiSMASH and BiG-SCAPE utilize gene-content evidence. Our Pfam baseline uses only the set of Pfam identifiers associated with each BGC. As a result, it does not include BiG-SCAPE’s duplication and adjacency terms. Another related work, BiG-SLiCE, is designed to compare families at large scale [3].

### Sequence-derived BGC representations

ESM-2 is a model trained on protein sequences that produces general sequence representations. Set Transformers use permutation-invariant attention mechanisms, like ISAB for differently sized sets [6]. Similarly, BGC-MAC / BGC-MAP utilize sequence representations for BGC classification and product matching, which are supervised prediction and cross-modal matching tasks, respectively [4]. The model predicts a BGC class or associates it with a molecular product representation. Our objective is different. Given a set of known producer BGCs, we rank other BGCs using BGC-side information and check if members of the same proxy group are retrieved. This distinction illustrates that strong performance in classification and product matching does not necessarily imply an advantage over explicit domain-content similarity for BGC-BGC retrieval.

## 3 Methods

### 3.1 Data and labels

Our dataset consists of 6,953 BGCs from 182 deduplicated *Streptomyces griseus* genomes. Of these, 5,325 have silver assignments to MIBiG reference groups [7]. We use them only to evaluate the retrieval system. They cannot be treated as evidence of a biosynthetic pathway or as direct product identities.

### 3.2 Group-disjoint split and retrieval protocol

Before model training, pair construction, or query generation, reference groups are split into training, validation, and test sets. This results in 4,211 training BGCs in 98 groups, 676 validation BGCs in 21 groups, and 438 test BGCs in 21 groups. The test groups are excluded from the entire pipeline, including Phase 1, Phase 2, validation-based score selection, and checkpoint selection. For each qualified test group four references are sampled, and only 16 test groups consist of more than 4 members.

For each group, we sample 100 deterministic query draws. We remove four sampled references from the candidate database, where remaining same-group members are positives. We mark unlabeled BGCs as unknown. For each query, all non-reference BGCs in the 438-BGC test set are ranked. Tied scores are handled using expected tie-aware metrics. The reported confidence intervals are 95% bootstrap intervals over the 16 eligible families.

### 3.3 Sequence-derived model

After processing genes through the ESM-2 model, we feed our sequence-derived model frozen 1,280-dimensional ESM-2 gene embeddings, normalized relative gene positions, and padding masks. Additional information, like product labels, activity labels, sequence-alignment statistics, and product-class indicators are excluded from model inputs. Pfam-augmented BGC-SetNet also receives a BGC-level Pfam inventory over a 2,218-token vocabulary built only from training BGCs, including padding and unknown tokens. The raw ESM baseline is calculated by normalizing the mean of the gene embeddings.

Our model, BGC-SetNet, is a Set Transformer with two induced self-attention blocks, 32 inducing points, and eight heads, followed by multihead attention pooling. In Phase 1, we mask 15% of gene embeddings. Let 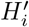 denote the contextual state of a masked gene. Then, the reconstruction before pooling is performed as

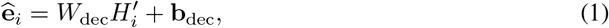

where the decoder maps to the frozen 1,280-dimensional ESM-2 space. Phase 1 uses training BGCs only. Phase 2 optimizes a family-aware contrastive objective on training groups. Five seeds are used: 20260810–20260814.

The validation-selected ensemble combines the learned Pfam-augmented BGC-SetNet cosine score and raw ESM cosine score. The selected BGC-SetNet + Pfam weights were 0.1, 0.1, 0.1, 0.2, and 0.4 across the five seeds; the test scores were never used to select these values.

### 3.4 Pfam and weighted-Pfam scores

For BGCs *A* and *B*, the unweighted baseline is

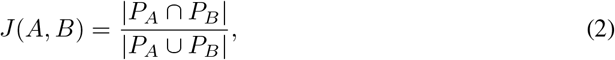

where *P*_*A*_ and *P*_*B*_ are their Pfam-identifier sets. The retrieval score is the maximum similarity between a candidate and the sampled references.

To test whether domain importance, rather than domain identity alone, could improve retrieval, we learn nonnegative domain weights *w*_*k*_ using only training-family labels. The differentiable score is

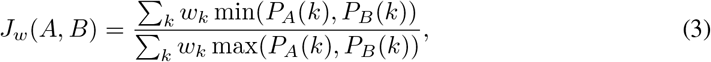

where *P*_*A*_(*k*) ∈ {0, 1} . The domain vocabulary is built from training BGCs only; validation and test-only domains map to an unknown token. Checkpoint selection uses validation Recall@50, and the final test score is direct weighted Jaccard without post-hoc tuning.

### 3.5 Statistical analysis

For each method, seed results are averaged within each held-out family before inference. Thus each paired test contains one observation per family, not family-by-seed, query, or BGC rows. We use an exact two-sided Wilcoxon signed-rank sign-flip enumeration after removing numerical zero differences. Comparisons are made against unweighted Pfam Jaccard. The 16-family effective sample size is reported explicitly; no seed flattening is used.

## 4 Results

### 4.1 Final held-out results

Table 1 reports the canonical five-seed aggregate. Pfam Jaccard and weighted Pfam are the strongest methods on the primary Recall@50 metric and are numerically indistinguishable. Pfam-augmented BGC-SetNet is lower than Pfam, while the ESM plus Pfam-augmented BGC-SetNet ensemble is close but lower.

**Table 1:** Final retrieval results under the group-disjoint split on 16 eligible held-out silver groups. Each cell shows the mean on the first line and the 95% family-bootstrap interval below it, after averaging five seeds within each family.

| Method | R@50 | MRR | MAP | nDCG@50 |
| --- | --- | --- | --- | --- |
| Raw ESM | 0.7946<br>[0.711, 0.870] | 0.2550<br>[0.171, 0.348] | 0.7251<br>[0.613, 0.837] | 0.8078<br>[0.724, 0.889] |
| BGC-SetNet + Pfam | 0.8472<br>[0.776, 0.912] | 0.2786<br>[0.180, 0.391] | 0.7771<br>[0.661, 0.884] | 0.8502<br>[0.772, 0.923] |
| Pfam Jaccard | 0.8788<br>[0.815, 0.938] | 0.3071<br>[0.195, 0.432] | 0.8480<br>[0.752, 0.934] | 0.9042<br>[0.846, 0.958] |
| ESM + BGC-SetNet<br>+ Pfam | 0.8769<br>[0.813, 0.936] | 0.3096<br>[0.197, 0.435] | 0.8503<br>[0.756, 0.934] | 0.9058<br>[0.849, 0.957] |
| Weighted Pfam | 0.8789<br>[0.814, 0.939] | 0.3069<br>[0.195, 0.436] | 0.8477<br>[0.749, 0.933] | 0.9040<br>[0.846, 0.958] |

**Table 2:** Exact paired family-level comparisons over 16 eligible test groups. Each observation is a five-seed family mean.

| Comparison | Metric | $\Delta$ | Wins/Ties/Losses | Raw $p$ | Holm $p$ |
| --- | --- | --- | --- | --- | --- |
| Ensemble vs. Pfam | Recall@50 | -0.00198 | 2/12/2 | 0.625 | 1.000 |
| Ensemble vs. Pfam | MRR | +0.00250 | 8/3/5 | 0.206 | — |
| Ensemble vs. Pfam | MAP | +0.00230 | 7/5/4 | 0.765 | — |
| Ensemble vs. Pfam | nDCG@50 | +0.00161 | 4/8/4 | 0.844 | — |
| Weighted Pfam vs. Pfam | Recall@50 | +0.00003 | 2/14/0 | 0.500 | 1.000 |

**Table 3:** Dataset and experimental configuration summary.

| Statistic | Value |
| --- | --- |
| Total atlas BGCs | 6,953 |
| Silver assigned BGCs | 5,325 |
| Train / validation / test BGCs | 4,211 / 676 / 438 |
| Train / validation / test groups | 98 / 21 / 21 |
| Eligible held-out groups | 16 |
| References per query | 4 |
| Query draws per group | 100 |

**Figure 1:**
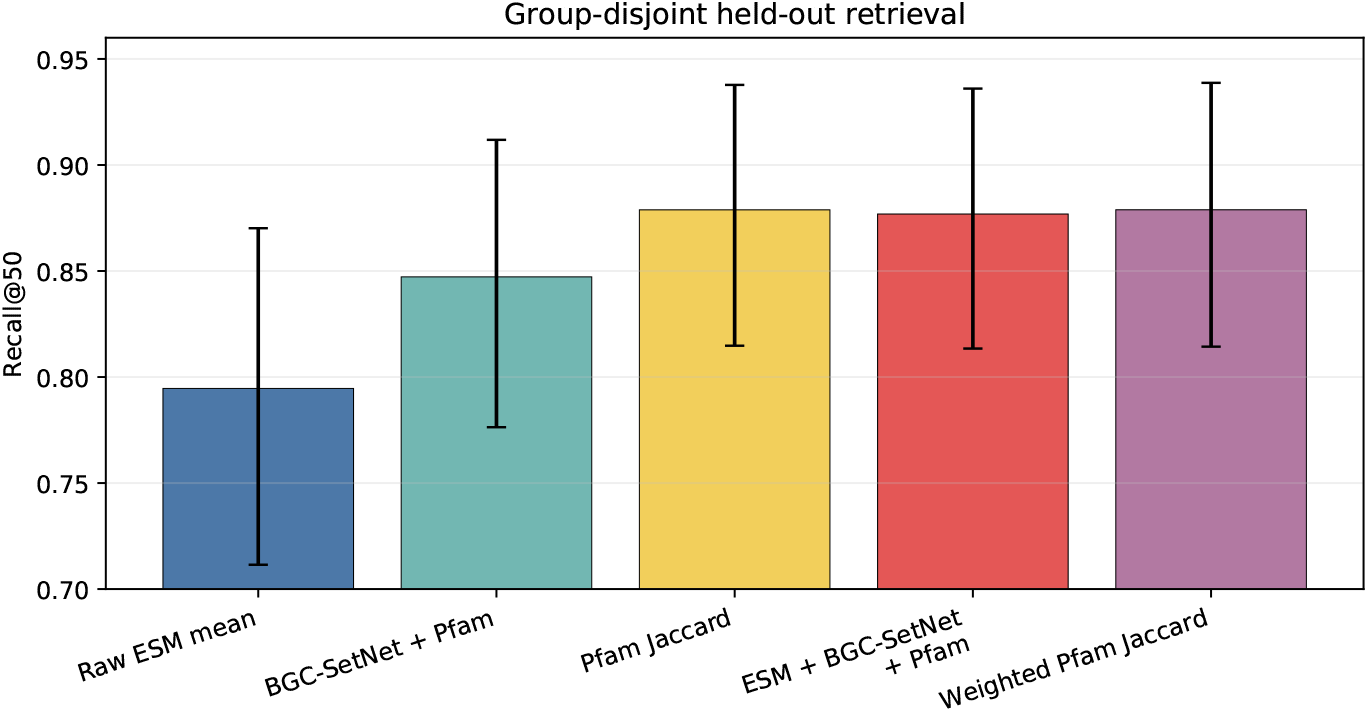
Final five-seed Recall@50 results with 95% family-bootstrap intervals. All values are generated from the canonical aggregate, not manually entered figure data.

**Figure 2:**
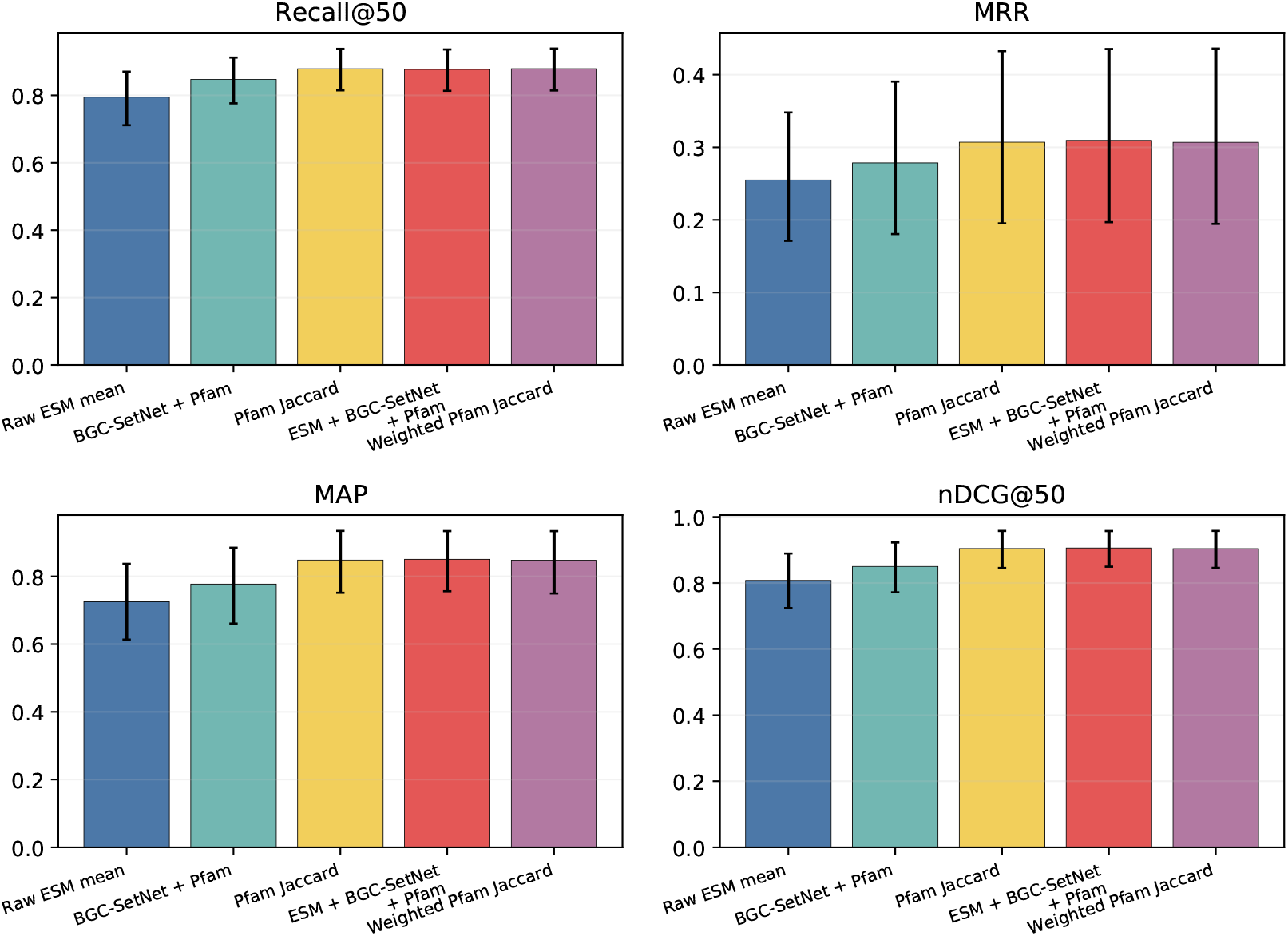
Final five-seed results across Recall@50, MRR, MAP, and nDCG@50. Explicit Pfam content remains the dominant signal; weighted Pfam Jaccard does not materially change the baseline.

### 4.2 Family-level comparisons

The ESM plus Pfam-augmented BGC-SetNet ensemble has a Recall@50 difference of − 0.00198 relative to Pfam Jaccard (exact *p* = 0.625; 2 wins, 12 ties, 2 losses). Weighted Pfam Jaccard has a difference of +0.00003 (exact *p* = 0.50; 2 wins, 14 ties, 0 losses). These are not meaningful improvements.

**Figure 3:**
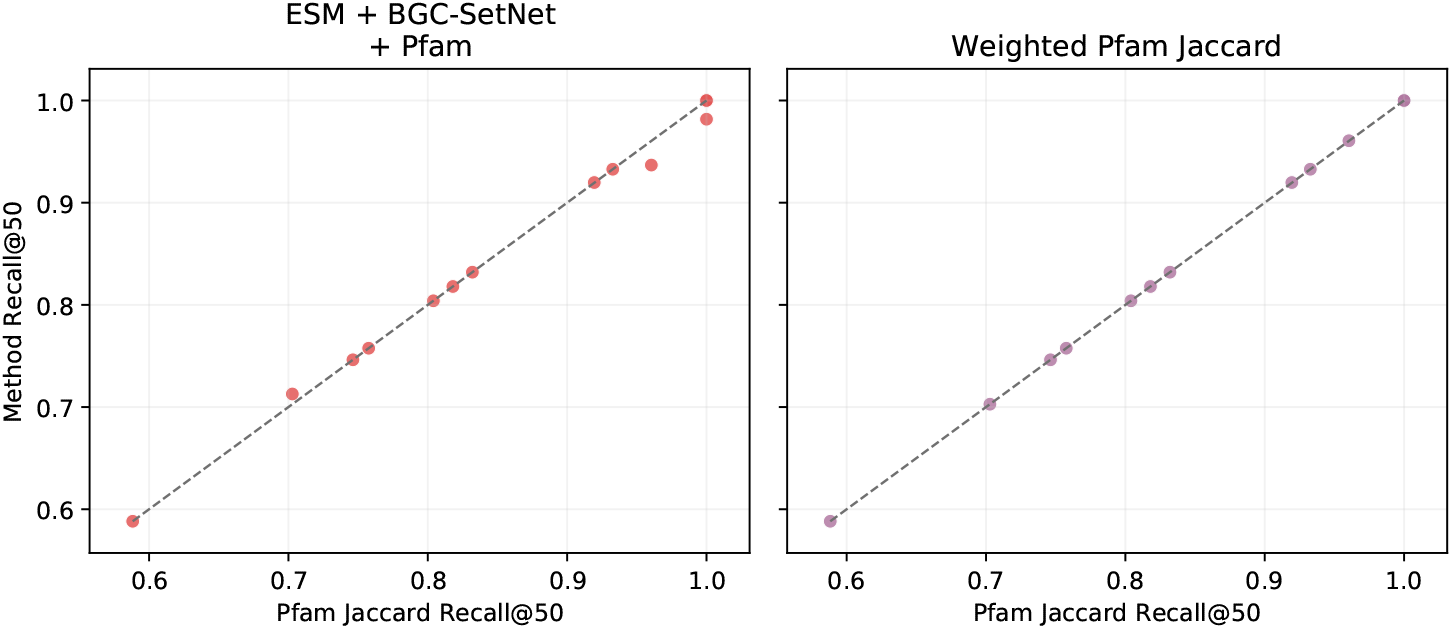
Per-family Recall@50 compared with Pfam Jaccard. The diagonal denotes parity. The learned ESM plus Pfam-augmented BGC-SetNet ensemble does not systematically exceed Pfam, and weighted Pfam is almost identical to the unweighted score.

**Figure 4:**
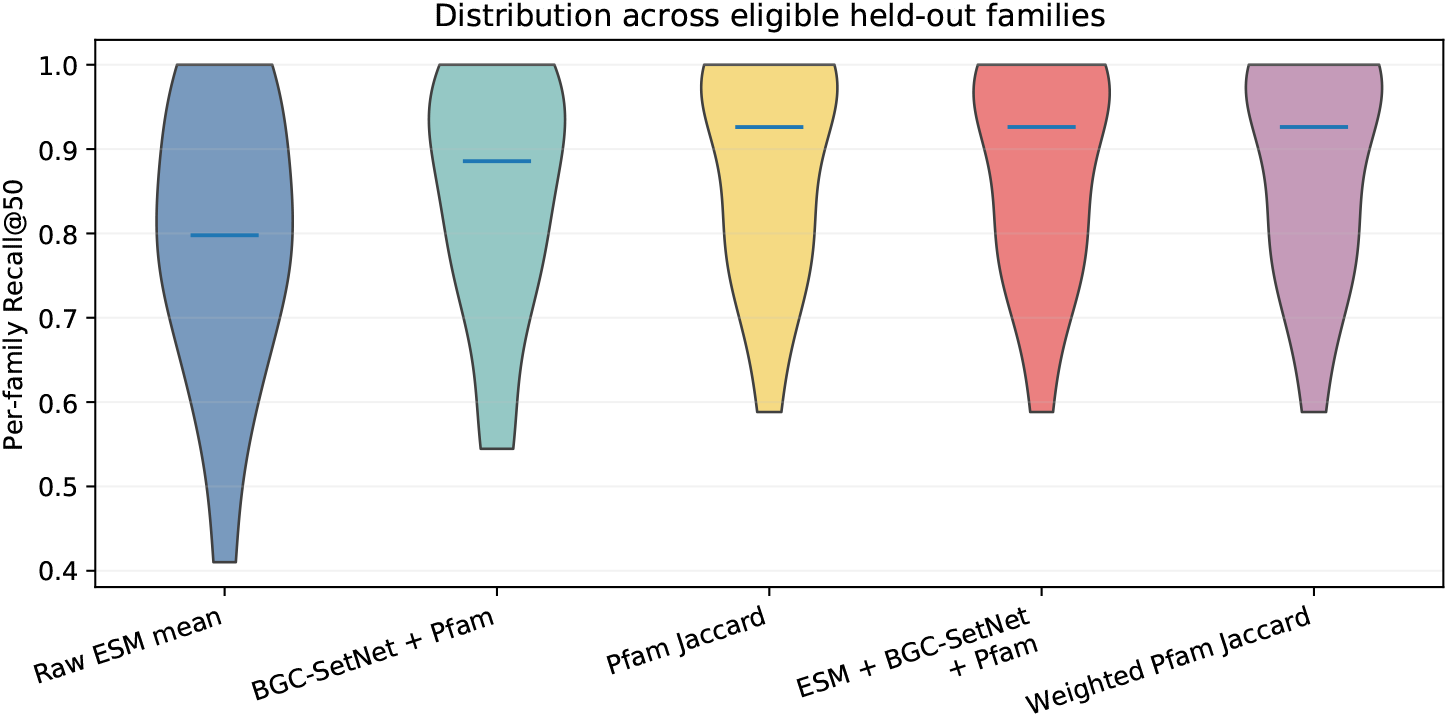
Distribution of family-level Recall@50 across the 16 eligible held-out groups. The plot uses family means after averaging five seeds; horizontal markers denote the median.

## 5 Alternative-pathway interpretation and limitations

Low Pfam Jaccard is an operational measure of divergent Pfam-domain content relative to the selected references. It does not establish a different biosynthetic pathway, a shared product, or a true alternative producer. The current benchmark is therefore suitable for candidate prioritization, not pathway discovery claims.

The principal limitations are silver labels rather than curated product identities, the missing machine-verifiable taxonomy manifest, and the small effective test size of 16 eligible families. The 16-family unit is a real power limitation: it supports transparent paired inference but makes small performance differences difficult to distinguish from sampling variation. We therefore report family-level effect sizes and confidence intervals and do not flatten seeds or query draws to increase the apparent sample size. A larger grouped cross-validation study would be useful, but it would require fold-specific retraining and validation and is outside the present fixed-holdout experiment. Because no independently adjudicated gold subset is available, we do not assign a numerical error rate to the silver labels; the reported metrics should be interpreted as performance against the proxy grouping used to construct this benchmark. The results also compare query-time scoring after Pfam and ESM preprocessing; they should not be described as a full antiSMASH runtime comparison. Finally, the weighted-Pfam model learns from the same domain inventory as the baseline, so its near-equivalence is diagnostic rather than evidence of independent biological validation.

## 6 Discussion

The central finding is not that sequence-derived representations are universally ineffective. It is that this controlled benchmark exposes a strong explicit-domain signal. Changing the aggregation architecture did not close the gap, adding the Pfam inventory to SetNet did not produce a meaningful primary-metric gain, and learning domain weights reproduced rather than improved standard Jaccard. The benchmark therefore provides a useful diagnostic for separating representation-learning gains from information already present in domain annotations.

This result clarifies what a stronger alternative-pathway study would require: curated product identities, a complete taxonomy and provenance manifest, pathway-level or gene-neighborhood features, and a new untouched grouped evaluation. Future models should be evaluated against the explicit-domain baseline rather than described as discovering alternative routes from low-overlap scores alone.

## 7 Conclusion

We present a reproducible five-seed evaluation framework for sequence-derived BGC retrieval. On 6,953 atlas BGCs and 16 eligible held-out silver groups, Pfam-based methods remain the strongest practical baselines. Neither the Pfam-augmented BGC-SetNet nor its validation-selected ESM plus Pfam-augmented BGC-SetNet ensemble provides a supported improvement, and a differentiable weighted-Pfam generalization is effectively identical to standard Pfam Jaccard. The correct biological interpretation is gene-content diversity relative to references, not proof of alternative biosynthetic pathways. The protocol and aggregate result artifacts provide a useful baseline for future work with curated labels and pathway-level evidence.

## Code and Data Availability

Manuscript source, the curated bioinformatics data-preparation pipeline, aggregate-analysis and figure-generation code, and compact family-level result artifacts are available at https://github.com/det3ctiv3/bgc_setnet. Final five-seed Pfam-augmented BGC-SetNet and weighted-Pfam checkpoints, per-seed configs, Pfam vocabularies, evaluation metadata, and a model card are available at https://huggingface.co/whiteh4t/bgc-setnet. Large raw genome files, generated atlas tables, and the frozen ESM-2 embedding matrix are not bundled in either source repository.

## A Dataset and experimental configuration

The final aggregate artifacts record split hash:

~~~
dc26fae17e54fd2ad41a9e10353b3da3
e0aacf3144b64f6ee62e8341b4360555
~~~

ESM plus Pfam-augmented BGC-SetNet runs use the two-phase configuration described above with five seeds. Weighted-Pfam runs use the same group split, a training-only 2,218-token Pfam vocabulary including padding and unknown tokens, direct differentiable weighted-Jaccard training, validation checkpoint selection, and direct test scoring.

